# Genetic deletion of NAPE-PLD induces context-dependent dysregulation of anxiety-like behaviors, stress responsiveness, and HPA-axis functionality in mice

**DOI:** 10.1101/2024.09.10.612324

**Authors:** Taylor J. Woodward, Diana Dimen, Emily Fender Sizemore, Sarah Stockman, Fezaan Kazi, Serge Luquet, Ken Mackie, Istvan Katona, Andrea G. Hohmann

## Abstract

The endocannabinoid (eCB) system regulates stress responsiveness and hypothalamic-pituitary-adrenal (HPA) axis activity. The enzyme *N*-acyl phosphatidylethanolamine phospholipase-D (NAPE-PLD) is primarily responsible for the synthesis of the endocannabinoid signaling molecule anandamide (AEA) and other structurally related lipid signaling molecules known as *N*-acylethanolamines (NAEs). However, little is known about how activity of this enzyme affects behavior. As AEA plays a regulatory role in stress adaptation, we hypothesized that reducing synthesis of AEA and other NAEs would dysregulate stress reactivity. To test this hypothesis, we evaluated wild type (WT) and NAPE-PLD knockout (KO) mice in behavioral assays that assess stress responsiveness and anxiety-like behavior. NAPE-PLD KO mice exhibited anxiety-like behaviors in the open field test and the light-dark box test after a period of single housing. NAPE-PLD KO mice exhibited a heightened freezing response to the testing environment that was further enhanced by exposure to 2,3,5-trimethyl-3-thiazoline (TMT) predator odor. NAPE-PLD KO mice exhibited an exaggerated freezing response at baseline but blunted response to TMT when compared to WT mice. NAPE-PLD KO mice also exhibited a context-dependent dysregulation of HPA axis in response to TMT in the paraventricular hypothalamic nucleus at a neuronal level, as measured by c-Fos immunohistochemstry. Male, but not female, NAPE-PLD knockout mice showed higher levels of circulating corticosterone relative to same-sex wildtype mice in response to TMT exposure, suggesting a sexually-dimorphic dysregulation of the HPA axis at the hormonal level. Together, these findings suggest the enzymatic activity of NAPE-PLD regulates emotional resilience and recovery from both acute and sustained stress.

**Significance Statement:** The endocannabinoid anandamide (AEA) regulates stress responsiveness and activity of the hypothalamic-pituitary-adrenal (HPA) axis. Currently, little is known about how an enzyme (i.e. *N*-acylphosphatidylethanolamine phospholipase-D (NAPE-PLD)) involved in the *synthesis* of AEA affects behavior. We hypothesized that genetic deletion of NAPE-PLD would dysregulate responsiveness to stress at a behavioral and neuronal level. Our studies provide insight into potential vulnerabilities to stress and anxiety that may result from dysregulation of the enzyme NAPE-PLD in people.

## Introduction

Cannabis has been reported to relieve feelings of stress and anxiety for millennia. However, the discovery of arachidonoyl ethanolamine (AEA), also known as anandamide, as an endogenous ligand for the cannabinoid type 1 (CB1) receptor opened the door for understanding the ways in which endocannabinoid (eCB) signaling regulates stress processing (Hillard et al., 2017). Stress can be described as an internal or external pressure, or disruption to homeostasis, which produces specific physiological and behavioral adaptations in an organism (Tan and Yip, 2018). While adaptive by nature, chronic or uncontrollable stress can be maladaptive to an organism without the proper coping resources. The eCB system modulates behavioral reactivity to environmental context in part via interactions with the hypothalamic-pituitary-adrenal (HPA) axis and depends on the state of the animal (Lutz et al., 2015).

The enzyme *N*-acyl phosphatidylethanolamine phospholipase-D (NAPE-PLD) catalyzes formation of lipids structurally related to AEA, known as *N*-acylethanolamines (NAEs). In rodents, NAPE-PLD is highly expressed in brain regions involved in stress processing and reward function, including the olfactory tubercle, the hippocampus and the hypothalamus (Egertová et al., 2008). In contrast to diacylglycerol lipase-alpha (DGL-a), the postsynaptic enzyme that produces the endocannabinoid 2-arachidonoylglycerol (2-AG), NAPE-PLD is predominantly restricted to *presynaptic* compartments in regions in which it is expressed (Nyilas et al., 2008; Pickel et al., 2012; Reguero et al., 2014). Although initially identified as a CB1 cannabinoid receptor agonist, AEA may also modulate neuronal activity via non CB1 mechanisms such as TRPV1 channels (Chávez et al., 2010; Grueter et al., 2010; Puente et al., 2011). Thus, disruption of AEA production may impact multiple physiological systems.

Little is known presently about how activity of NAPE-PLD impacts behavior. In the present study, we asked how global genetic deletion of NAPE-PLD would influence reactivity to stress and HPA axis functionality. To this end, we examined the performance of WT and NAPE-PLD KO (Liu et al., 2008) mice of both sexes in preclinical behavioral tests that assess anxiety-like behaviors and stress responsiveness. To evaluate a response to a mild stressor, we compared genotypic differences of singly-housed mice to mice that were group-housed. To investigate whether deletion of the NAPE-PLD gene alters neuronal activity in brain regions where NAPE-PLD is highly expressed, we performed immunolabeling in the hippocampus and hypothalamus against the protein product of the immediate early gene *c-fos*, which is a widely used and reliable marker for neuronal activity (Anisimova et al., 2023). Finally, we observed the behavioral response of WT and NAPE-PLD KO mice to 2,4,5-trimethyl-3-thiazoline (TMT), a component of fox feces and urine, which serves as an ethologically relevant uncontrollable stressor (Janitzky et al., 2015; Rosen et al., 2015). We also evaluated HPA axis response in NAPE-PLD KO and WT mice of both sexes by quantifying c-Fos protein expression in the hypothalamic paraventricular nucleus (PVN) as well as levels of circulating corticosterone at baseline and in response to stressors of increasing intensity.

## Methods

### Animals

All procedures were approved by the Institutional Animal Care and Use Committee at Indiana University Bloomington. NAPE-PLD KO mice on a C57BL/6J background mice were bred as described previously (Leishman et al., 2016). NAPE-PLD KO mice in this study were generated either from a heterozygous cross (+/-with +/-) or as the F1 generation of a non-sibling homozygous pair generated from the heterozygous cross. WT mice were littermates or were supplemented with C57BL/6J from Jackson Laboratories (Bar Harbor, ME, USA). Mice were single-or group-housed (2-4 per cage, separated by sex) depending on the experimental conditions required. All mice had ad libitum access to water and food and maintained on 12-hour light/dark cycle (lights on at 8:00 AM). Unless specifically mentioned, all mice were between 12 and 16 weeks of age at time of testing.

### Chemicals

2,3,5-trimethyl-3-thiazoline (TMT) was obtained from BioSRQ (Sarasota, FL).

### Behavioral Assays

Prior to all behavioral experiments, mice were handled by the experimenter in their colony room. All behavioral experiments and tissue collection were performed in the animals’ light cycle between 11:00 am and 6:00 pm. Group-housed mice were marked on the tail with a permanent marker 1-2 days prior to behavioral testing for identification.

#### Open Field Test

Mice were brought into the room and habituated in their home cage on a table for 30-90 minutes prior to testing. Mice were placed in activity meters (Omnitech Superflex Nodes, Omnitech, Columbus, Ohio) and were automatically recorded by photobeams interpreted by Fusion 6.5 software starting the moment they entered the arena as described previously (Murphy et al., 2017; Iyer et al., 2024). An elevated set of sensors recorded vertical activity time, or the time mice spent in exploratory rearing positions. The chambers were illuminated with tungsten bulbs at ∼80 lux, and a white noise generator provided a steady sound level of 62-63 dB in the arenas. Animal behavior was recorded for 30 minutes, after which mice were promptly removed from each arena. Arenas were thoroughly cleaned with 70% ethanol in between animals.

#### Light-Dark Box

Animals were brought into the testing room and habituated in their home cage on a table for at least 30 minutes prior to testing. Mice were placed in activity meters (Omnitech Superflex Nodes) containing a light-dark box insert and were automatically recorded by Fusion 6.5 software starting the moment they entered the arena as described previously (Dvorakova et al., 2021) . The chambers were illuminated with tungsten bulbs at 400-450 lux (dark side <3 lux), and a white noise generator provided a steady sound level of 62-63 dB in the arenas. Activity in the light-dark box was measured for 5 minutes, after which the animals were promptly removed.

#### Elevated Plus Maze

The elevated plus maze was performed as described previously (Murphy et al., 2017). Mice were brought into the testing room and habituated in their cages on a table for at least 30 minutes prior to testing. Mice were placed in the elevated plus maze facing the open arm away from the experimenter. Activity on the EPM was recorded with an overhead camera for 5 minutes, after which the mouse was returned to its home cage and the EPM was cleaned with 70% EtOH. Fecal boli left on the EPM were counted after the animal was returned to its cage. Illumination in the open arms was ∼60 lux and ∼15 lux in the closed arms. A white noise generator provided a steady sound level of 62-63 dB in the arenas. Mice were then returned to their housing room. Behavior was analyzed offline with ezTrack open-source software (Pennington et al., 2019).

#### Forced Swim Test

The forced swim test was performed as described previously (Maciel et al., 2022). Mice were brought to a testing room containing a cylinder of water maintained at 25° C. Mice were placed into the cylinder for 6 minutes and video recorded for the duration of the test. After 6 minutes, mice were removed from the cylinder of water, dried with a towel, and returned to their home cage. Immobility time during the last 4 minutes was scored manually offline by an experimenter blinded to treatment.

#### Marble Burying

Marble burying was carried out as described previously by our group (Slivicki et al., 2018). Mice were single housed for 3 days prior to the first behavioral procedure. The same cohort of mice went through marble burying followed by nestlet shredding. A standard polycarbonate cage was filled with 1 inch of level saw dust bedding. Then, 20 standard glass marbles were placed in a 4x5 grid. The mouse was placed in a corner of the cage away from the marbles and allowed to interact with the marbles for 30 minutes. Then, the mouse was removed, and an image of the cage captured digitally. Marbles were considered covered if they were 2/3 or greater concealed in bedding. The number of marbles covered was counted after 30 minutes.

#### Nestlet Shredding

Nestlet shredding was carried out as described previously by our group (Slivicki et al., 2018). After the marble burying assay, a cotton nestlet square was added to the home cage for 2 days prior to nestlet shredding assay. On the day of the assay, mice were placed in a clean holding cage before any bedding material, food, and water bottle were removed from the home cage. The cotton nestlet was weighed and cut into 6 even pieces and placed equidistantly in the cage. Mice were returned to the home cage and allowed to interact with the nestlets for 100 minutes before removal. An image of the cage was taken and the number of cleared zones of cage (out of 6) was recorded. The shredded portion of nestlet was removed from the intact nestlet and intact nestlet was weighed. Percent shredded was determined by the amount of nestlet shredded during the assay divided by its original weight x 100.

#### Predator Odor Exposure

To control for social buffering/social amplification of stress, mice were single-housed for several days prior to odor exposure experiments. Odor exposures were performed in acrylic chambers (4.5 x 5 x 11 inches) and videos of behavior were recorded using Logitech C240 webcams and Logitech Capture software. Mice in their cages were brought to a holding room near the fume hood at least 30 minutes prior to habituation/exposure sessions. On habituation day (Day 0), mice were placed in the experimental chamber (which was situated in the fume hood), videorecorded for 20 minutes, and placed back in their home cage, after which they were returned to their colony room. Then, 24 hours later, mice underwent a similar procedure, except that a 20 mL liquid scintillation vial containing a single gauze square was taped in the top of the chamber to allow passive diffusion of odor. Water, butyric acid (Sigma-Aldrich, St. Louis, MO) or 2,4,5-trimethyl-3-thiazoline (TMT; BioSRQ, Florida, USA) were pipetted at volumes of 1 to 25 µL directly onto the gauze pad in the scintillation vial immediately prior to placing mice in the chamber. The ambient sound level in the fume hood was ∼77dB and the lights were dimmed to 100 lux.

To minimize odor contamination, water-exposed mice were tested in a chamber that had never been used for TMT exposures and water-exposed mice were tested each exposure day prior to TMT-exposed mice. In between each odor exposure, chambers were thoroughly cleaned at least twice with ethanol (or until the experimenter was unable to detect any traces of TMT). Great care was taken to minimize odor contamination prior to TMT exposure, and all materials that came into contact with TMT (gloves, paper towels, pipette tips) were stored in a closed container within the fume hood until the completion of the day’s experiments. Videos were analyzed offline using the FreezeAnalysis module of ezTrack open-source software (Pennington et al., 2019, 2021). Behavioral videos were calibrated to videos taken of an empty chamber as per instruction of ezTrack designers. A mouse was counted as freezing if it performed no other behaviors beyond breathing for a period of more than 1 second. Odor exposures were performed in batches of two, with males and females tested jointly, each in their separate experimental chambers.

#### Corticosterone Quantification

CORT levels were compared between behaviorally naïve, group-housed mice (low stress controls) WT and NAPE-PLD KO, or immediately after the 20-minute exposure to either water (10 µL) or TMT (10 µL; an intense stressor) to capture the peak of CORT release. The TMT exposure condition was used as the intense stressor, whereas the water exposure condition, which also involved placement in the fume hood, represented a mild stressor. Cardiac blood was collected from the right atrium. Samples were allowed to coagulate at room temperature for 60 minutes, after which they were centrifuged at 1300 g for 15 minutes (4°C) and serum was stored at −80 °C until analysis. Samples were analyzed with an enzyme-linked immunosorbent (ELISA) assay kit purchased from Enzo Life Sciences (CAT: ADI-900-097, New York, USA). Per manufacturer’s instructions, serum samples were diluted 1:8 and absorbance fit to a standard curve to determine sample concentration.

### Neuronal activation in WT and NAPE-PLD KO mice following exposure to stimuli with different levels of aversiveness

Behaviorally naïve group housed WT and NAPE-PLD KO mice were transcardially perfused for the investigation of the baseline activity level in the hippocampus and PVN. Mice that were exposed to water or TMT were returned to their home cage and left undisturbed until they were perfused (95 minutes after the onset of odor exposure) to capture the peak of c-Fos protein expression (Janitzky et al., 2015).

#### Sample preparation

Mice were transcardially perfused with saline (20 mL) followed by ice-cold 4% paraformaldehyde (PFA) solution (100 mL) prepared in phosphate buffer (0.1 M PB). The brains were dissected and postfixed in the same fixative solution overnight at 4°C, then transferred to the PB containing sodium-azide (0.05%). Coronal brain sections (50 μm thick) were prepared using a Leica VT 1200S vibratome. Sections were collected in PB containing 0.05% sodium-azide and stored at 4°C.

#### Fluorescent immunolabeling

We used c-Fos immunohistochemistry to both reveal whether there is a difference between the baseline activity level of neurons in the hippocampus and the PVN, and investigate neuronal activity changes following specific stimuli in each genotype. We labeled the c-Fos immediate-early gene protein product to quantify the activity of hippocampal CA3 and DG neurons, as well as the neuronal activity in the PVN. After washing with PB several times, free floating brain sections were blocked in 5% bovine serum albumin (BSA) in PB containing 0.9% NaCl and Triton X-100 (TPBS) for 1 hour at room temperature and then incubated with the primary c-Fos antibody (Abcam, Boston, MA, catalog number: ab190289) in 2.5% BSA-TPBS for overnight at a 1:2000 dilution. After incubation, sections were washed with PBS four times for 10 minutes and then incubated in the anti-rabbit A594 secondary antibody (Abcam, Boston, MA, catalog number: ab150080) in 2.5% BSA-TPBS with DAPI for 4 hours at room temperature. After several washing steps with PB, the sections were mounted onto a glass slide and cover-slipped in VECTASHIELD Antifade Mounting Medium.

#### Microscopy and image analysis

Confocal images were acquired using Nikon A1 Confocal Microscope equipped with epi-and fluorescent illumination (Lumencor, SPECTRA X light engine). Images were captured at 522 X 522 pixel resolution using 4-20x objectives at an optical thickness of 5 μm. c-Fos-positive neurons were counted automatically using QuPath open-source software (Bankhead et al., 2017). Quantification performed by QuPath was verified manually in 2 brain sections from each mouse at 2 planes using standardized settings (contrast, intensity) across the sections. Full resolution of the images was maintained until the final versions, which were adjusted to a resolution of 300 dpi. Fluorescent intensity of c-Fos positive cells was quantified by taking the average pixel intensity of a 3-image z-stack taken 5 µm above and below the focus point automatically detected by the microscope.

#### Experimental Design and Statistics

Sample group size was determined by power analyses conducted in pilot studies. Behavioral data were analyzed by One-way, or Two-way Analysis of Variance (ANOVA) as appropriate followed by a Bonferroni post-hoc test for multiple comparisons. Immunohistochemical data was analyzed using the Mann-Whitney test, Two-way ANOVA, or Spearman’s correlation, as appropriate. A full table of statistical analyses used for all figures is given in Table 1. GraphPad Prism version 10.0 (GraphPad Software, La Jolla, CA, USA) was used for analyses and graph creation. Adobe Photoshop 23.5 was used to display IHC images. Data are presented as mean ± SEM. In all instances, p<0.05 was considered statistically significant.

**Table 1.** Statistics.

| Figure | Test | Interaction (2 way ANOVA) | Pvalue | Sex | Pvalue | Genotype | Pvalue |
| --- | --- | --- | --- | --- | --- | --- | --- |
| 1A | 2 way ANOVA | Interaction: F (1, 64) = 1.220 | P=0.2735 | Sex: F (1, 64) = 4.343 | P=0.0412 | Genotype: F (1, 64) = 0.2589 | P=0.6126 |
| 1B | 2 way ANOVA | Interaction: F (1, 64) = 22.80 | P<0.0001 | Sex: F (1, 64) = 15.48 | P=0.0002 | Genotype: F (1, 64) = 30.32 | P<0.0001 |
| 1C | 2 way ANOVA | Interaction: F (1, 64) = 1.505 | P=0.2243 | Sex: F (1, 64) = 4.293 | P=0.0423 | Genotype: F (1, 64) = 2.943 | P=0.0911 |
| 1D | 2 way ANOVA | Interaction: F (1, 36) = 1.075 | P=0.3068 | Sex: F (1, 36) = 1.159 | P=0.2888 | Genotype: F (1, 36) = 11.85 | P=0.0015 |
| 1E | 2 way ANOVA | Interaction: F (1, 36) = 0.1088 | P=0.7434 | Sex: F (1, 36) = 2.262 | P=0.1413 | Genotype: F (1, 36) = 19.06 | P=0.0001 |
| 1F | 2 way ANOVA | Interaction: F (1, 36) = 1.072 | P=0.3073 | Sex: F (1, 36) = 5.576 | P=0.0237 | Genotype: F (1, 36) = 4.056 | P=0.0515 |
| 1G | 2 way ANOVA | Interaction: F (1, 75) = 3.081 | P=0.0833 | Sex: F (1, 75) = 0.008684 | P=0.9260 | Genotype: F (1, 75) = 35.78 | P<0.0001 |
| 1H | 2 way ANOVA | Interaction: F (1, 55) = 2.453 | P=0.1231 | Sex: F (1, 55) = 1.031 | P=0.3143 | Genotype: F (1, 55) = 16.76 | P=0.0001 |
| 1I | 2 way ANOVA | Interaction: F (1, 75) = 2.016 | P=0.1598 | Sex: F (1, 75) = 47.89 | P<0.0001 | Genotype: F (1, 75) = 3.391 | P=0.0695 |
| 2A | 2 way ANOVA | Interaction: F (1, 31) = 0.3499 | P=0.5585 | Sex: F (1, 31) = 3.164 | P=0.0851 | Genotype: F (1, 31) = 1.610 | P=0.2140 |
| 2B | 2 way ANOVA | Interaction: F (1, 72) = 4.003 | P=0.0492 | Sex: F (1, 72) = 1.144 | P=0.2883 | Genotype: F (1, 72) = 18.15 | P<0.0001 |
| 2C | 2 way ANOVA | Interaction: F (1, 49) = 1.989 | P=0.1647 | Sex: F (1, 49) = 19.08 | P<0.0001 | Genotype: F (1, 49) = 3.120 | P=0.0836 |
| 2D | 2 way ANOVA | Interaction: F (1, 49) = 0.2688 | P=0.6065 | Sex: F (1, 49) = 8.886e-005 | P=0.9925 | Genotype: F (1, 49) = 9.386 | P=0.0035 |
| 2E | 2 way ANOVA | Interaction: F (1, 43) = 4.439 | P=0.0410 | Sex: F (1, 43) = 0.8956 | P=0.3493 | Genotype: F (1, 43) = 1.311 | P=0.2585 |
| 2F | 2 way ANOVA | Interaction: F (1, 33) = 0.0005155 | P=0.9820 | Sex: F (1, 33) = 1.634 | P=0.2100 | Genotype: F (1, 33) = 0.1845 | P=0.6703 |
| 2G | 2 way ANOVA | Interaction: F (1, 33) = 0.4290 | P=0.5170 | Sex: F (1, 33) = 1.088 | P=0.3045 | Genotype: F (1, 33) = 2.335 | P=0.1360 |
| 3E | Spearman's Correlation | Spearman r = .9069 | P<0.0001 |  |  |  |  |
| 3F | 2 way ANOVA | Interaction: F (1, 12) = 2.736 | P=0.1240 | Sex: F (1, 12) = 8.571 | P=0.0127 | Genotype: F (1, 12) = 33.90 | P<0.0001 |
| 3G | 2 way ANOVA | Interaction: F (1, 12) = 17.82 | P=0.0012 | Sex: F (1, 12) = 68.06 | P<0.0001 | Genotype: F (1, 12) = 145.4 | P<0.0001 |
| 3H | Mann-Whitney test | n/a | P=0.0001 |  |  |  |  |
| 3I | Mann-Whitney test | n/a | P<0.0001 |  |  |  |  |
| 3J | 2 way ANOVA | Interaction: F (1, 12) = 0.01773 | P=0.8963 | Sex: F (1, 12) = 15.57 | P=0.0019 | Genotype: F (1, 12) = 0.1276 | P=0.7271 |
| 3K | Mann-Whitney test | n/a | P=0.0007 |  |  |  |  |
| 3L | Mann-Whitney test | n/a | P=0.5160 |  |  |  |  |
| 3M | 2 way ANOVA | Interaction: F (1, 12) = 2.267 | P=0.1580 | F (1, 12) = 2.327 | P=0.1531 | F (1, 12) = 2.327 | P=0.1531 |
| 4B | 2 way ANOVA | Interaction: F (76, 627) = 9.901 | P<0.0001 | Time: F (8.107, 267.5) = 31.99 | P<0.0001 | Odor: F (4, 33) = 46.43 | P<0.0001 |
| 4C | One way ANOVA | F (4, 33) = 46.43 | P<0.0001 |  |  |  |  |
| 4D | Mixed effects | Interaction: F (2, 51) = 2.846 | P=0.0673 | Context: F (2, 51) = 400.8 | P<0.0001 | Sex: F (1, 53) = 1.132 | P=0.2921 |
| 4E | Mixed Effects | Interaction: F (2, 45) = 13.25 | P<0.0001 | Context: F (2, 45) = 111.3 | P<0.0002 | Sex: F (1, 47) = 0.04095 | P=0.8405 |
| 4F | 2 way ANOVA | Interaction: F (1, 100) = 2.470 | P=0.1192 | Sex: F (1, 100) = 7.113 | P=0.0089 | Genotype: F (1, 100) = 34.57 | P<0.0001 |
| 4G | 2 way ANOVA | Interaction: F (1, 46) = 0.4986 | P=0.4837 | Sex: F (1, 46) = 0.05699 | P=0.8124 | Genotype: F (1, 46) = 13.63 | P=0.0006 |
| 4H | 2 way ANOVA | Interaction: F (1, 50) = 0.4166 | P=0.5216 | Sex: F (1, 50) = 5.312 | P=0.0254 | Genotype: F (1, 50) = 8.284 | P=0.0059 |
| 5G | 2 way ANOVA | Interaction: F (2, 18) = 1.113 | P=0.3501 | Context: F (2, 18) = 228.1 | P<0.0001 | Sex: F (1, 18) = 5.149 | P=0.0358 |
| 5H | 2 way ANOVA | Interaction: F (2, 18) = 0.05634 | P=0.9454 | Context: F (2, 18) = 77.90 | P<0.0001 | Sex: F (1, 18) = 0.06259 | P=0.8053 |
| 5I | 2 way ANOVA | Interaction: F (2, 18) = 39.76 | P<0.0001 | Context: F (2, 18) = 289.4 | P<0.0001 | Genotype: F (1, 18) = 0.1047 | P=0.7500 |
| 5J | 2 way ANOVA | Interaction: F (2, 18) = 14.75 | P=0.0002 | Context: F (2, 18) = 78.56 | P<0.0001 | Genotype: F (1, 18) = 2.330 | P=0.1443 |
| 5K | Spearman's Correlation | Spearman r: 0.7500 | P=0.0012 |  |  |  |  |
| 5L | Spearman's Correlation | Spearman r: 0.6441 | P=0.0085 |  |  |  |  |
| 6B | 2 way ANOVA | Interaction: F (1, 30) = 2.651 | P=0.1139 | Sex: F (1, 30) = 2.804 | P=0.1044 | Genotype: F (1, 30) = 6.635 | P=0.0152 |
| 6C | 2 way ANOVA | Interaction: F (2, 44) = 1.038 | P=0.3628 | Context: F (2, 44) = 22.04 | P<0.0001 | Sex: F (1, 44) = 2.291 | P=0.1373 |
| 6D | 2 way ANOVA | Interaction: F (2, 37) = 1.830 | P=0.1746 | Context: F (2, 37) = 28.66 | P<0.0001 | Sex: F (1, 37) = 0.1269 | P=0.7237 |
| 6E | 2 way ANOVA | Interaction: F (2, 40) = 0.8350 | P=0.4413 | Context: F (2, 40) = 19.06 | P<0.0001 | Genotype: F (1, 40) = 3.082 | P=0.0868 |
| 6F | 2 way ANOVA | Interaction: F (2, 41) = 5.200 | P=0.0097 | Context: F (2, 41) = 36.76 | P<0.0001 | Genotype: F (1, 41) = 18.32 | P=0.0001 |

## Results

### NAPE-PLD KO mice exhibited increased avoidance of the center of the open field after single housing

To examine how environmental and developmental factors influence the way in which NAPE-PLD regulates anxiety-like behavior, we assessed the open field performance of 12-16-week-old WT and NAPE-PLD KO mice that were group-housed or single-housed, as well as aged (35+ weeks old) single-housed mice of each genotype. Group-housed NAPE-PLD KO mice did not differ in center time from WT controls, although males spent slightly more time in the center than female mice regardless of genotype (Fig 1A) [Interaction: F (1, 64) = 1.220, P=0.2735; Sex: F (1, 64) = 4.343, P=0.0412; Genotype: F (1, 64) = 0.2589, P=0.6126]. Group-housed NAPE-PLD KO spent less time rearing than WT mice, females showed less time rearing than males overall, and group-housed male, but not female, NAPE-PLD KO mice spent less time in exploratory rearing behaviors than WT controls (Fig 1B) [Interaction: F (1, 64) = 22.80, P<0.0001; Sex: F (1, 64) = 15.48, P=0.0002; Genotype: F (1, 64) = 30.32, P<0.0001]. NAPE-PLD KO mice did not differ from WT controls in locomotor activity during the assay (Fig 1C) [Interaction: F (1, 64) = 1.505, P=0.2243; Sex: F (1, 64) = 4.293, P=0.0423; Genotype: F (1, 64) = 2.943, P=0.0911]. After spending 3 weeks single-housed, a separate cohort of NAPE-PLD KO mice spent less time in the center of the arena than WT controls irrespective of sex (Fig 1D) [Interaction: F (1, 36) = 1.075, P=0.3068; Sex: F (1, 36) = 1.159, P=0.2888; Genotype: F (1, 36) = 11.85, P=0.0015]. Single-housed NAPE-PLD KO mice of both sexes (12-16 weeks old) spent less time in exploratory rearing behaviors than WT controls (Fig 1E) [Interaction: F (1, 36) = 0.1088, P=0.7434; Sex: F (1, 36) = 2.262, P=0.1413; Genotype: F (1, 36) = 19.06, P=0.0001]. Single-housed males traveled less than females overall, irrespective of genotype, and single housed NAPE-PLD KO mice trended towards a modest decrease in distance traveled relative to their WT counterparts that approached statistical significance (Fig. 1F) [Interaction: F (1, 36) = 1.072, P=0.3073; Sex: F (1, 36) = 5.576, P=0.0237; Genotype: F (1, 36) = 4.056, P=0.0515). When both single-housed and tested at an advanced age (35+ weeks old), NAPE-PLD KO mice of both sexes spent less time in the center of the arena than age-matched WT controls under the same conditions (Fig 1G) [Interaction: F (1, 75) = 3.081, P=0.0833; Sex: F (1, 75) = 0.008684, P=0.9260; Genotype: F (1, 75) = 35.78, P<0.0001]. Similarly, aged (35+ weeks old) NAPE-PLD KO mice spent less time in exploratory rearing behaviors than age-matched WT controls, although the main effect of sex was not observed in the aged animals (Fig 1H) [Interaction: F (1, 55) = 2.453, P=0.1231; Sex: F (1, 55) = 1.031, P=0.3143; Genotype: F (1, 55) = 16.76, P=0.0001]. Aged NAPE-PLD KO mice also trended towards a modest decrease in distance traveled that did not reach statistical significance (Fig. 1I) [Interaction: F (1, 75) = 2.016, P=0.1598; Sex: F (1, 75) = 47.89, P<0.0001; Genotype: F (1, 75) = 3.391, P=0.0695]. Regardless of genotype, we found that female mice traveled longer distances than males at all ages and irrespective of housing conditions (Fig 1C, F, I). Heat maps show mean locomotor activity in group-housed (Fig. 1J) and single-housed (Fig. 1K) WT and NAPE-PLD KO mice.

**Figure 1:**
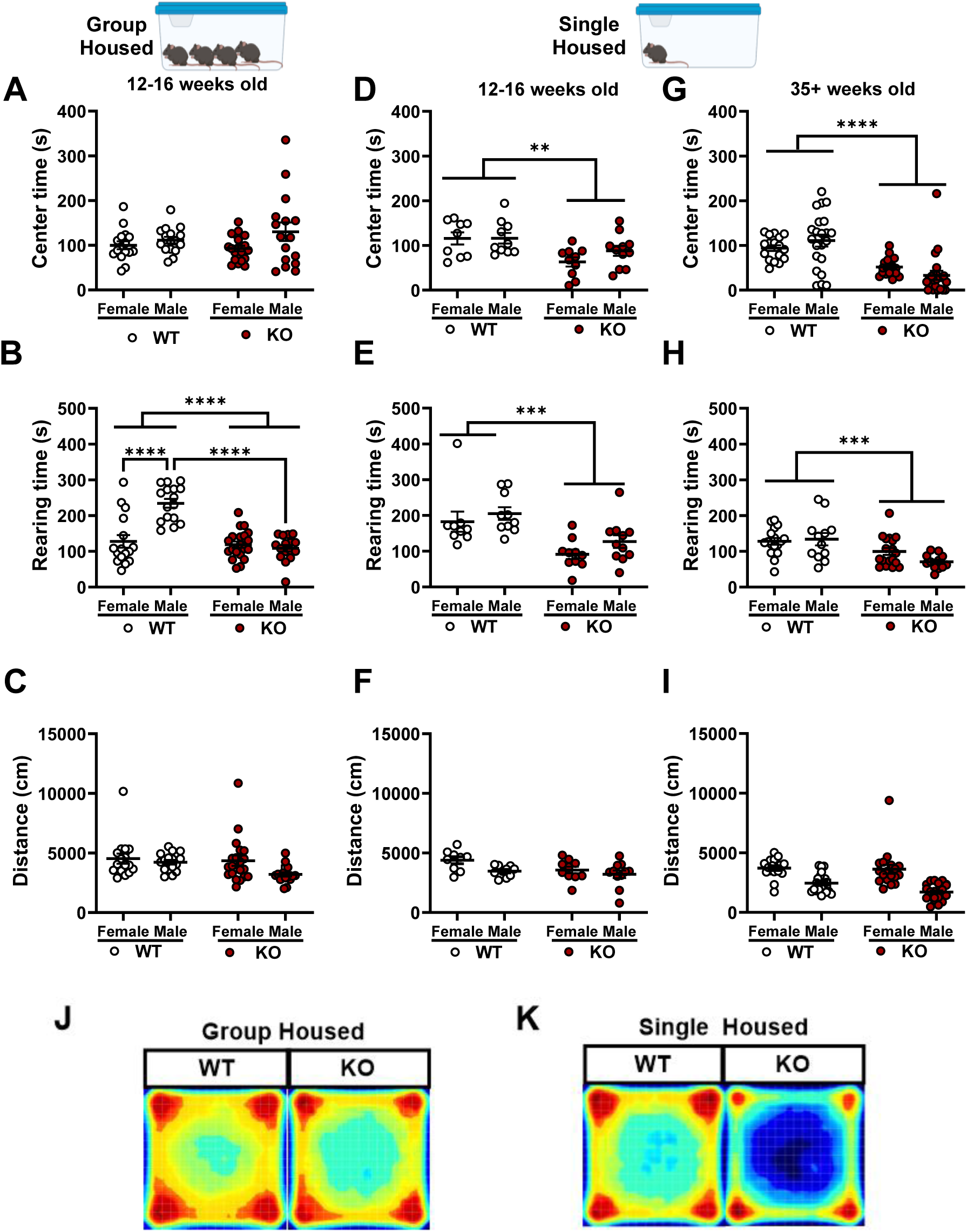
Single housed, but not group housed, NAPE-PLD KO mice exhibited an anxiety-like behavioral phenotype compared to WT controls. In the open field test, 12-16 week old group-housed NAPE-PLD KO mice did not differ from group-housed WT controls in the amount of time spent in the center of the arena **(A)**. In group-housed mice, male WT mice spent more time than both female WT and male NAPE-PLD KO mice (Sex x Genotype Interaction: p<0.0001) while no sex differences were observed in NAPE-PLD KO mice **(B).** Group-housed NAPE-PLD KO mice did not differ from WT controls in distance traveled **(C)**. When single-housed for 3 weeks prior to the test, 12-16 week old NAPE-PLD KO mice spent less time in the center of the arena than WT controls **(D)**. 12-16 weeks old NAPE-PLD KO mice spent less time in exploratory rearing behaviors than WT controls without a significant sex by genotype interaction **(E)**. Single housed NAPE-PLD KO mice did not differ from single housed WT controls in distance traveled **(F)**. When aged past 35 weeks, single housed NAPE-PLD KO mice that were spent less time in the center of the arena than age-matched WT controls **(G)** and less time in exploratory rearing behaviors **(H)**. Aged NAPE-PLD KO mice did not differ from WT controls in distance traveled **(I).** In all age groups and housing conditions, female mice traveled longer distances than male mice regardless of genotype in all age groups and housing conditions (Sex: p<0.05) **(G, H, I)**. Representative group heatmaps are presented for a cohort of group housed 12-16 weeks old WT and NAPE-PLD KO mice **(J)** as well as a cohort of single housed 35+ week old WT and NAPE-PLD KO mice **(K)**. Red regions indicate higher time spent in an area, while blue regions indicate lower amount of time spent in an area. Data are presented as the mean ± S.E.M. Multiple comparisons from two-way ANOVA were followed by a Bonferroni post-hoc test. ****p <.0001, ***p <.001, **p <.01, *p <.05 WT vs NAPE-PLD KO. Sample size: n = 8-22 mice per sex per genotype.

### Context-dependent dysregulation of emotional behaviors and stress adaptation in NAPE-PLD KO mice

Genotypic differences were not observed in the number of times that group-housed 12-16 week old mice entered the light side of the arena (Fig 2A) in the light-dark box [Interaction: F (1, 31) = 0.3499, P=0.5585; Sex: F (1, 31) = 3.164, P=0.0851; Genotype: F (1, 31) = 1.610, P=0.2140]. However, aged single-housed NAPE-PLD KO mice (especially males) entered the light side of the arena fewer times than WT controls (Fig 2B) [Interaction: F (1, 72) = 4.003, P=0.0492; Sex: F (1, 72) = 1.144, P=0.2883; Genotype: F (1, 72) = 18.15, P<0.0001]. In the elevated plus maze, females spent more time on the open arm than males regardless of genotype, although no reliable genotypic differences were present in open arm time (Fig 2C) [Interaction: F (1, 49) = 1.989, P=0.1647; Sex: F (1, 49) = 19.08, P<0.0001; Genotype: F (1, 49) = 3.120, P=0.0836]. However, while WT controls rarely defecated during the 5-minute test, NAPE-PLD KO mice left more fecal boli on the elevated plus maze compared to WT mice (Fig 2D) [Interaction: F (1, 49) = 0.2688, P=0.6065; Sex: F (1, 49) = 8.886e-005, P=0.9925; Genotype: F (1, 49) = 9.386, P=0.0035]. In the forced swim test, male NAPE-PLD KO mice spent more time immobile than male WT controls, while no genotypic difference was present among females (Fig 2E) [Interaction: F (1, 43) = 4.439, P=0.0410; Sex: F (1, 43) = 0.8956, P=0.3493; Genotype: F (1, 43) = 1.311, P=0.2585]. When tested in their home cage (i.e. under less aversive conditions), single-housed NAPE-PLD KO mice did not differ from WT controls in nestlet shredding (Fig 2F) or marble burying (Fig 2G) behaviors [Fig. 2F, Nestlet Shredding: Interaction: F (1, 33) = 0.0005155, P=0.9820; Sex: F (1, 33) = 1.634, P=0.2100; Genotype: F (1, 33) = 0.1845, P=0.6703. Fig. 2G, Marble Burying: Interaction: F (1, 33) = 0.4290, P=0.5170; Sex: F (1, 33) = 1.088, P=0.3045; Genotype: F (1, 33) = 2.335, P=0.1360].

**Figure 2:**
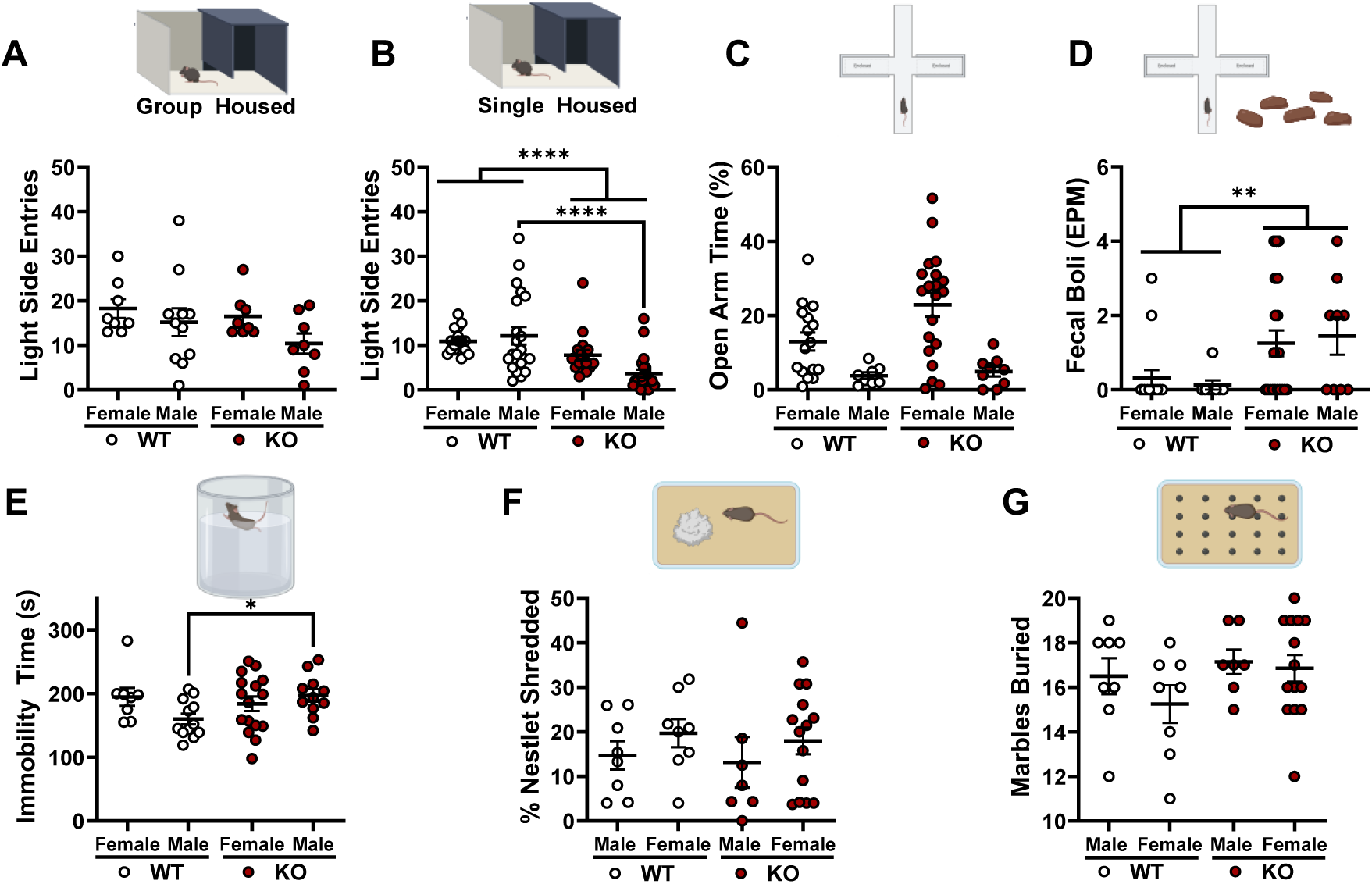
Dysregulation of emotional behaviors and stress adaption in NAPE-PLD KO mice is dependent upon assay aversiveness and context. When group housed, 12-16 week old NAPE-PLD KO mice did not differ in number of light side entries from WT controls in the light-dark box assay **(A).** However, when aged and single housed, NAPE-PLD KO mice entered the light side of the box fewer times than age-matched WT controls, a genotypic effect that was especially prevalent in males **(B)**. In the elevated plus maze assay, male mice spent less time in the open arm than female mice, but no reliable genotypic differences were observed **(C)**. However, NAPE-PLD KO mice of both sexes defecated a greater number of fecal pellets than WT controls during testing in the elevated plus maze **(D)**. In the forced swim test, male NAPE-PLD KO mice spent more time immobile than male WT controls, while no genotypic difference was observed in females **(E)**. Single housed 12–16-week-old NAPE-PLD KO mice did not differ from WT controls in performance in home-cage assays, the nestlet shredding test **(F)** and the marble burying test **(G)**. Data is presented as the mean ± S.E.M. Multiple comparisons from two-way ANOVA were followed by a Bonferroni post-hoc test: ****p<0.0001, * p<0.05 for shown comparisons. Sample size: n=8-18 per sex per genotype.

### Genetic deletion of NAPE-PLD reduced c-Fos immunolabeling in the hippocampus

To investigate how deletion of NAPE-PLD would affect basal neuronal activity in the hippocampus (where it is highly expressed), we performed immunolabeling against the protein product of the immediate early gene *c-fos*. Representative images are shown for WT mice of both sexes (Fig 3A, B) as well as NAPE-PLD KO mice of both sexes (Fig 3C, D). Automatic cell counts generated by QuPath were independently verified by manual counting to ensure accuracy. Cell counts generated by QuPath correlated strongly with manual cell counts (Fig 3E) [Spearman r = .9069, P<0.0001]. While c-Fos+ cell counts were higher on average in males than females in the CA3 (Fig 3F) and dentate gyrus (DG) (Fig 3G) subregions of the hippocampus, NAPE-PLD KO mice, irrespective of sex, exhibited reduced numbers of c-Fos+ cells compared to WT controls in both regions and the interaction was significant in the DG [Fig. 3F CA3: Interaction: F (1, 12) = 2.736, P=0.1240; Sex: F (1, 12) = 8.571, P=0.0127; Genotype: F (1, 12) = 33.90, P<0.0001. Fig. 3G, DG: Interaction: F (1, 12) = 17.82, P=0.0012; Sex: F (1, 12) = 68.06, P<0.0001; Genotype: F (1, 12) = 145.4, P<0.0001]. To determine whether the lower number of active neurons in NAPE-PLD KO mice is associated with an overall reduction in c-Fos expression in these cells, we additionally measured the mean fluorescent intensity (FI, each cell’s average pixel intensity from a z-stack) of all c-Fos+ cells in WT and NAPE-PLD KO mice for each brain region. In CA3, mean fluorescent intensity of c-Fos+ cells analyzed from female (Fig 3H) and male (Fig 3I) NAPE-PLD KO mice was on average lower than WT c-Fos+ mean cell fluorescent intensity [Females: p<0.0001, Mann-Whitney Test; Males: p<0.0001, Mann-Whitney Test]. In the DG, a similar decrease in mean c-Fos+ cell fluorescent intensity was present in female (Fig 3L) but not male (Fig 3M) NAPE-PLD KO mice [Females: p=0.0007, Mann-Whitney Test; Males: p<0.5160, Mann-Whitney Test]. Interestingly, the ratio of High FI/Low FI c-Fos+ cells was reduced in the CA3 region (Fig 3J) but not the DG of NAPE-PLD KO mice (Fig 3M) [CA3 Region: Interaction: F (1, 12) = 0.01773, P=0.8963; Sex: F (1, 12) = 0.1276, P=0.7271; Genotype: F (1, 12) = 15.57, P=0.0019. DG Region: Interaction F (1, 12) = 2.267, P=0.1580; Sex: F (1, 12) = 2.327, P=0.1531; Genotype: F (1, 12) = 2.327, P=0.1531]

**Figure 3.**
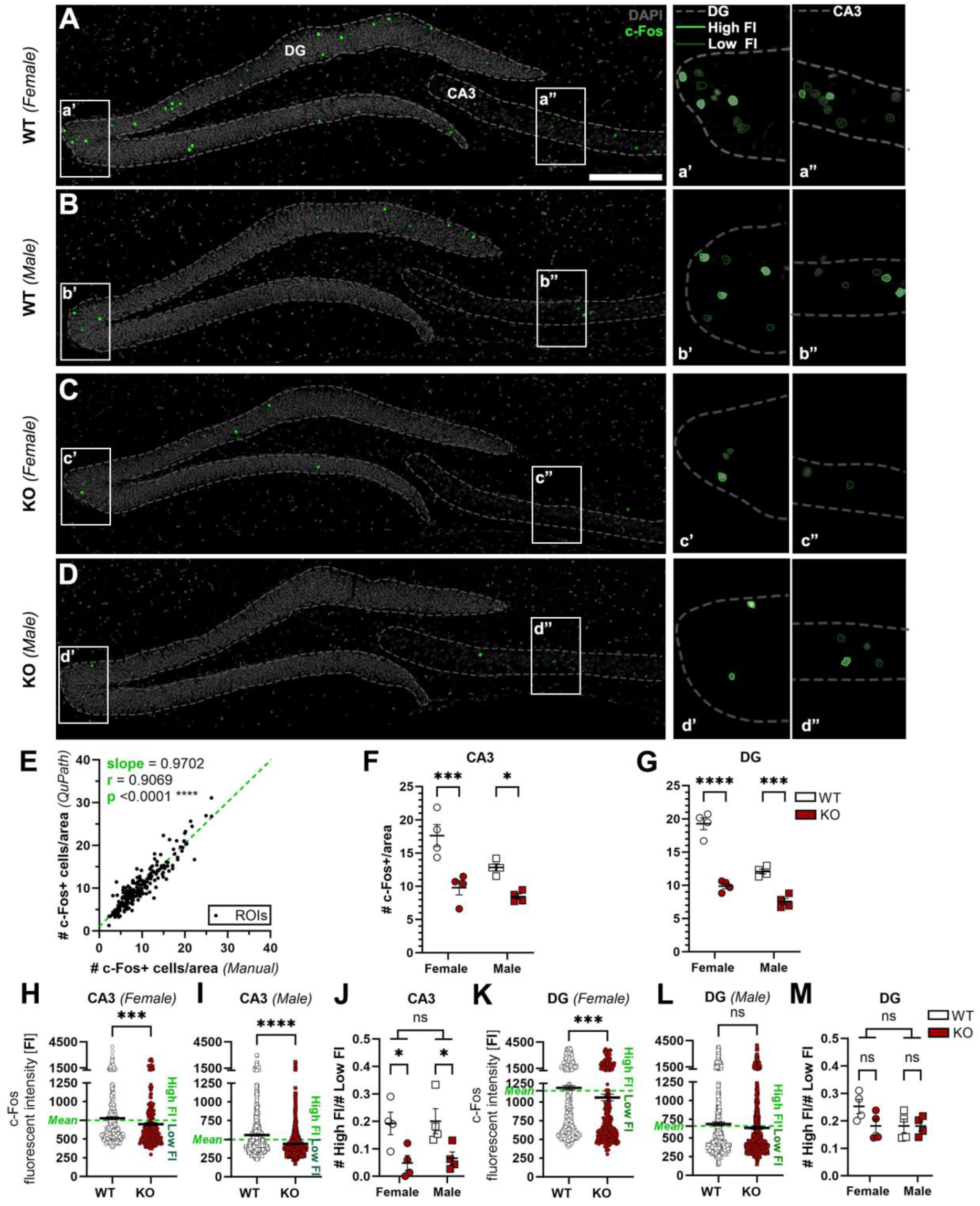
NAPE-PLD KO mice exhibit reductions in basal hippocampal c-Fos positive cell counts and intensity. (**A-D**) Representative confocal micrographs of c-Fos expression in the hippocampus from experimentally naïve female (**A**) and male WT **(B)** and female (**C**) and male NAPE-PLD KO **(D)** mice. Dashed lines mark the border of the dentate gyrus (DG) and the CA3 region. Inserts demonstrate cells marked as low fluorescent intensity (Low FI) and high fluorescent intensity (High FI) labeled by dark green line and light green border, respectively. Scale bar: 200 µm. **(E)** Verification of the deep-learning-based cell counting analysis by correlating the c-Fos+ cell numbers counted by QuPath program with manual analysis, with each dot representing one ROI that was analyzed (Spearman correlation; r = 0.9069, p < 0.0001****). c-Fos-positive (+) cell counts were lower in NAPE-PLD KO mice of both sexes in both the CA3 **(F)** and in the DG **(G)** regions. In F and G, each dot represents one animal. Mean fluorescent intensity of all c-Fos-positive cells located in the CA3 region was lower on average in NAPE-PLD KO mice of both sexes **(H, I)** compared to WT controls (each dot represents one c-Fos-positive cell). Within each sex, the average intensity of all measured c-Fos-positive cells (both WT and KO grouped together) is labeled on the Y-axis (Mean) with a green dashed line **(H, I, K, and L)**. Cells above the dashed line were considered as High FI cells, while cells below the dashed line were considered as Low FI cells. In the CA3 region, NAPE-PLD KO mice of both sexes exhibited a reduction in the ratio of #High FI/#Low FI cell counts for each animal **(J,** each data point represents the ratio of one animal). Compared to WT cell fluorescent intensity, a reduction in mean fluorescent intensity of c-Fos-positive cells located in the DG was observed in female **(K)** but not male **(L)** NAPE-PLD KO mice (each dot represents one c-Fos+ cell). Interestingly, the ratio of High FI/Low FI was not reduced in the DG of NAPE-PLD KO mice **(M,** each dot represents one animal) Each area was normalized to 50,000 µm^2^ for presentation. Data are presented as the mean ± S.E.M. Multiple comparisons from two-way ANOVA followed by Bonferroni post-hoc test: ****p<0.0001, ***p<0.001, **p<0.01, * p<0.05 for shown comparisons. Sample size: n=4 per sex per genotype.

### NAPE-PLD KO mice exhibited increased freezing in the absence of odor and heightened freezing evoked by predator odor is blunted in comparison to WT mice

As NAPE-PLD KO mice responded differently than WT to a mild ambient stressor (single-housing), we asked whether NAPE-PLD KO mice would respond differently to an acute intense stressor. We selected exposure to TMT as an ethologically relevant stressor known to activate the HPA axis (Janitzky et al., 2015). As seen in the experimental schematic (Fig 4A), mice were habituated to the testing environment (acrylic chamber placed in a fume hood) on day 1 and exposed to odor 24 hours later (Fig 4A). In initial pilot experiments performed in male WT mice, TMT at quantities of 1, 10, and 25 µL (pipetted onto the gauze square) elicited a quantity-dependent freezing response, whereas control odors water and butyric acid (10 µL) did not elicit a freezing response in WT mice under the same conditions (Fig 4B) [Interaction: F (76, 627) = 9.901, P<0.0001; Time: F (8.107, 267.5) = 31.99, P<0.0001; Odor: F (4, 33) = 46.43, P<0.0001]. Analysis of the area under the curve of total freezing time from the 20-minute exposure confirmed that TMT produced concentration dependent increases in total freezing time compared to the water exposed control group (Fig 4C) [One way ANOVA: F (4, 33) = 46.43, P<0.0001; p <0.01 for each comparison]. Subsequent genotypic comparisons were performed in response to 10 µL of water or TMT, as this volume of TMT elicited a freezing response that plateaued at about 50%, enabling us to detect subtle behavioral changes without ceiling or floor effects. WT mice spent more time freezing in response to TMT exposure than in response to water exposure or during habituation. (Fig 4D) [Mixed effects model Interaction: Interaction: F (2, 51) = 2.846, P=0.0673; Context: F (2, 51) = 400.8, P<0.0001; Sex: F (1, 53) = 1.132, P=0.2921]. NAPE-PLD KO mice spent more time freezing in response to TMT exposure than during habituation or water exposure irrespective of sex (Fig 4E) [Mixed effects model: Interaction: F (2, 45) = 13.25, P<0.0001; Context: F (2, 45) = 111.3, P<0.0002; Sex: F (1, 47) = 0.04095, P=0.8405]. When compared directly to WT controls, NAPE-PLD KO mice spent more time freezing during habituation (Fig 4F), and females spent more time freezing than males [Interaction: F (1, 100) = 2.470, P=0.1192; Sex: F (1, 100) = 7.113, P=0.0089; Genotype: F (1, 100) = 34.57, P<0.0001]. In response to water exposure, NAPE-PLD KO mice similarly spent more time freezing than WT controls (Fig 4G) [Interaction: F (1, 46) = 0.4986, P=0.4837; Sex: F (1, 46) = 0.05699, P=0.8124; Genotype: F (1, 46) = 13.63, P=0.0006]. Conversely, NAPE-PLD KO mice that were exposed to TMT spent less time freezing than WT controls, and overall females spent less time freezing than males (Fig 5H) [Interaction: F (1, 50) = 0.4166, P=0.5216; Sex: F (1, 50) = 5.312, P=0.0254; Genotype: F (1, 50) = 8.284, P=0.0059].

**Figure 4:**
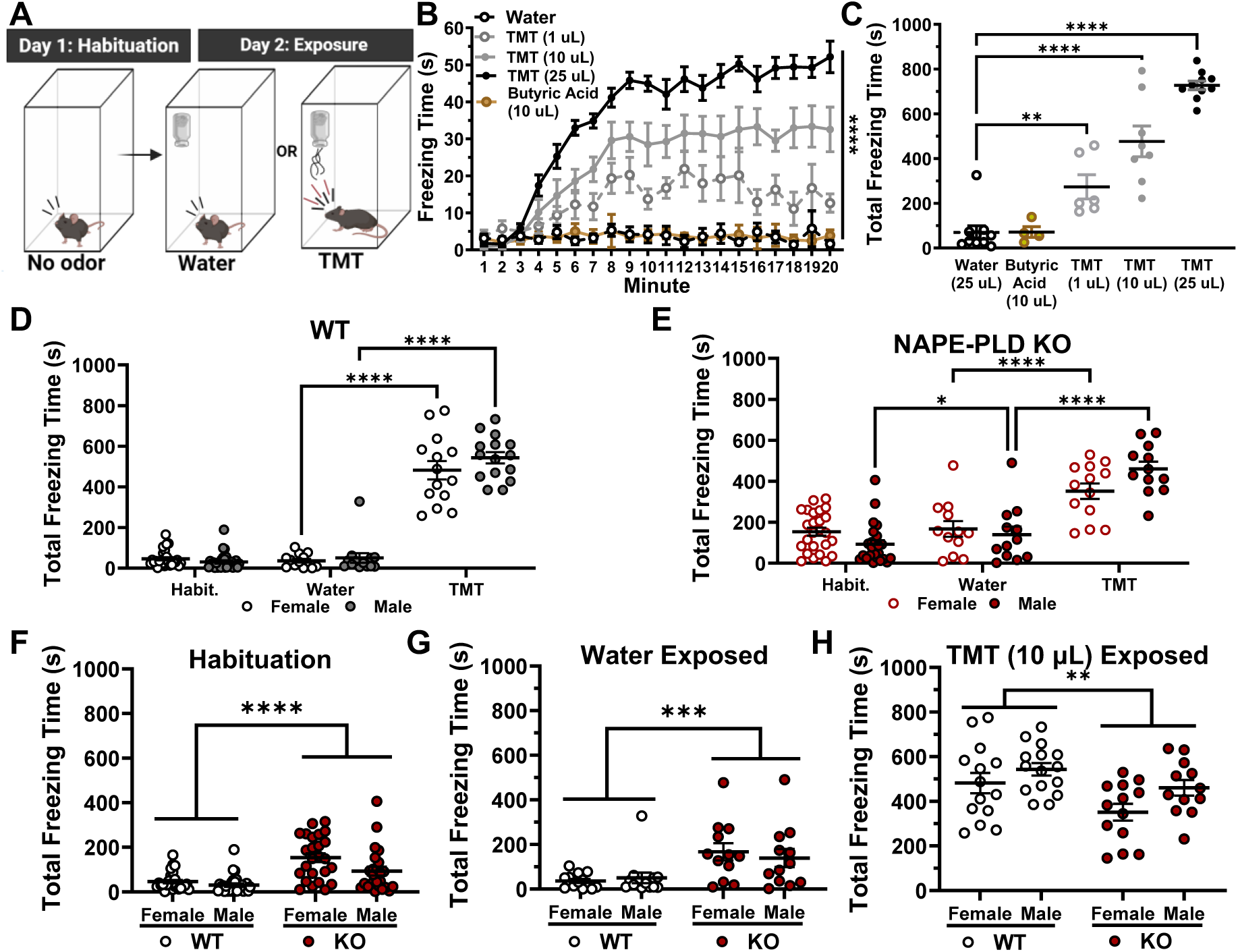
NAPE-PLD KO mice exhibited enhanced freezing in the absence of predator odor and a blunted freezing response to predator odor stress. A schematic shows the experimental protocol **(A)**. TMT (1,10, and 25 μL), but not the control odor butyric acid (10 μL), induced a quantity-dependent and time dependent freezing response that generally plateaued around 8 minutes after exposure initiation (**B**). TMT exposure produced concentration dependent increases in total freezing time compared to water exposure (in the fume hood) whereas butyric acid failed to do so **(C)**. WT mice of either sex showed increased total freezing time (measured as area under the curve, during the 20-minute test) in response to TMT, but not water, exposure **(D)**. NAPE-PLD KO mice showed more total freezing in response to TMT than in response to water exposure, irrespective of sex. Response to water exposure and habituation was similar in NAPE-PLD KO mice (**E**). Upon habituation to the testing environment (**F**) and in response to water exposure (**G**), KO mice of either sex spent more total time freezing than did WT mice. Conversely, NAPE-PLD KO mice that were exposed to TMT spent less time freezing than TMT-exposed WT mice **(H)**. Data are presented as the mean ± S.E.M. Multiple comparisons from one-way ANOVA (C), Linear Mixed Effects Model (D and E), and Two-Way ANOVA (F, G and H) followed by Bonferroni post-hoc test: ****p<0.0001, ***p<0.001, **p<0.01, for shown comparisons. Sample size: n=12-16 per sex per genotype per odor exposure group.

**Figure 5.**
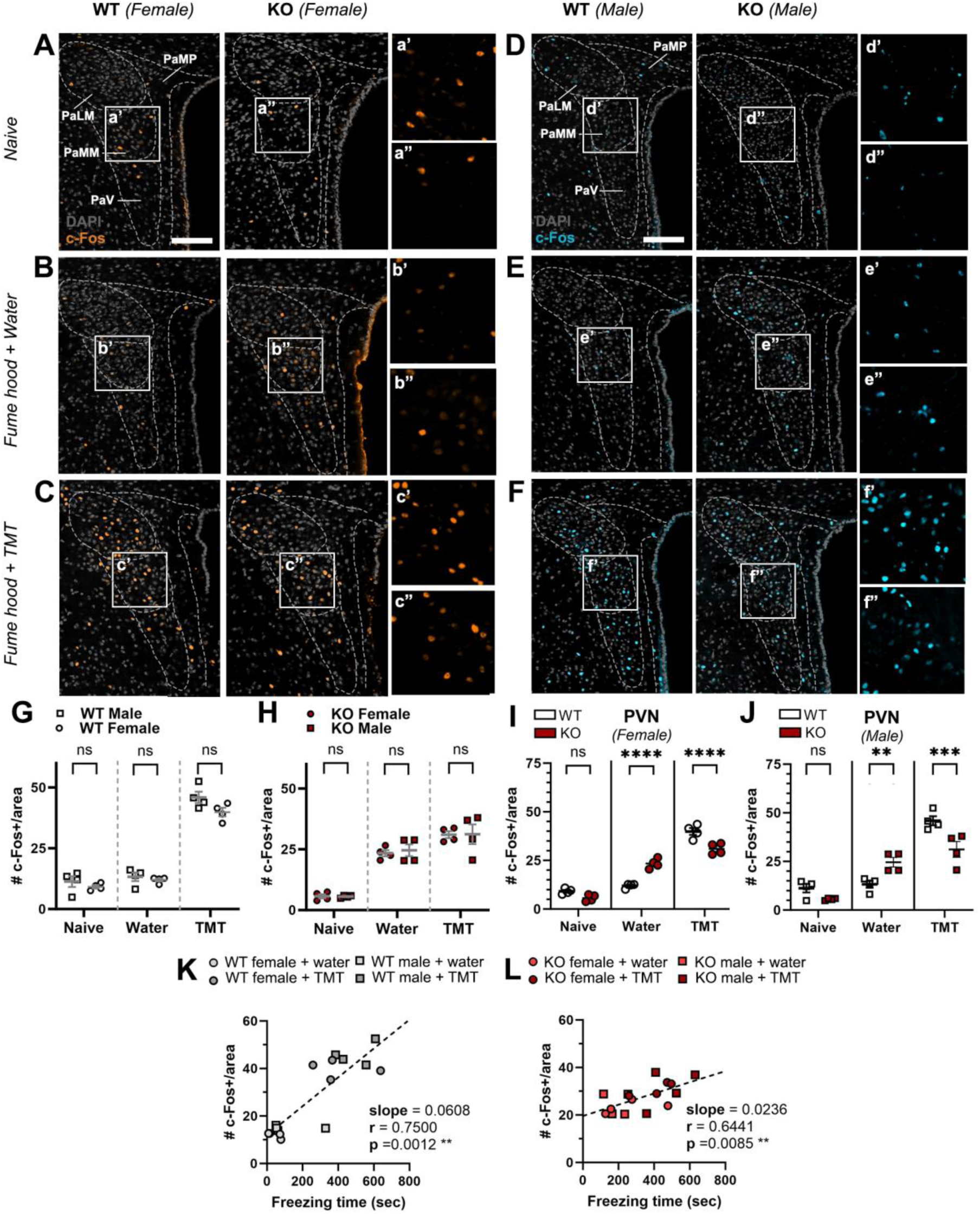
Expression of c-Fos in the hypothalamus was amplified in NAPE-PLD KO mice in response to testing conditions and not further increased by TMT exposure. Representative confocal micrographs demonstrating the c-Fos expression in behaviorally naïve **(A, D)**, water-exposed **(B, E)** and TMT-exposed (10 µL) **(C, F)** mice of both sexes (female data in orange, left two columns; male data in blue, right two columns). Dashed lines mark the border of the subregions of the hypothalamic paraventricular nucleus (PVN), including the paraventricular lateral magnocellular (PaLM), medial magnocellular (PaMM), medial parvocellular (PaMP) and the ventral periventricular (PaV) subregions. Inserts demonstrate the changes in the number of the c-Fos-positive cells across treatment conditions and between the genotypes (Scale bars: 100 µm). In WT mice, TMT exposure increased the number of c-Fos+ cells in PVN compared to water-exposed and naïve conditions **(G)**. Compared with naïve controls, NAPE-PLD KO mice exhibited an amplified response in c-Fos+ cell quantities in the water-exposed group (i.e. in response to the testing environment), which was not further increased in TMT-exposed NAPE-PLD KO groups **(H)**. Within-sex genotypic comparisons illustrate that c-Fos+ cell counts were elevated in both female **(I)** and male **(J)** NAPE-PLD KO mice exposed to water, while c-Fos+ cell counts were lower in TMT-exposed NAPE-PLD KO mice compared to WT controls. Freezing time and c-Fos+ cell counts correlated with freezing times in WT **(K)** and NAPE-PLD KO **(L)** mice, though the slope was flatter among NAPE-PLD KO mice due to the amplified freezing and c-Fos expression in the water-exposed group. Data are presented as the mean ± S.E.M. Multiple comparisons from two-way ANOVA followed by Bonferroni post-hoc test: ****p<0.0001, ***p<0.001, **p<0.01, * p<0.05 for shown comparisons. Each dot represents one animal. Sample size n=4 mice per sex per genotype.

### Expression of c-Fos in the hypothalamus was amplified in NAPE-PLD KO mice in response to testing conditions and not further increased by TMT exposure

To determine whether neuronal activity patterns are associated with the observed behavioral phenotype, a subset of WT and NAPE-PLD KO mice were perfused 95 minutes after the initiation of odor exposure for c-Fos immunolabeling. Behaviorally naïve subjects were also tested for c-Fos response to control for neural response to the experimental setting (i.e., placement in the experimental chamber within the fume hood can be expected to involve some level of stress). We quantified the number of c-Fos+ cell in the hypothalamic paraventricular nucleus (PVN), in female (orange, left two columns) and male (blue, right two columns) mice of both genotypes (Fig 5A-F). Representative images are show for each treatment group in experimentally naïve mice (Fig 5A, D), in water-exposed mice (Fig 5B, E), and in mice exposed to 10 µL of TMT (Fig 5C, F). In WT mice, TMT exposure elevated c-Fos+ cell count compared to water-exposed and naïve controls of both sexes, and males had higher quantities of c-Fos+ cells across contexts (Fig 5G) [Interaction: F (2, 18) = 1.113, P=0.3501; Context: F (2, 18) = 228.1, P<0.0001; Sex: F (1, 18) = 5.149, P=0.0358]. In NAPE-PLD KO mice, exposure to water (i.e. testing conditions) elevated c-Fos+ cell quantities, and TMT exposure did not further increase c-Fos+ cell counts (Fig 5H). Unlike WT mice, no sex differences were present across contexts in NAPE-PLD KO mice [Interaction: F (2, 18) = 0.05634, P=0.9454; Context: F (2, 18) = 77.90, P<0.0001; Sex: F (1, 18) = 0.06259, P=0.8053]. This effect (amplified response from testing environment that was not further increased by TMT exposure) was observed in both female (Fig 5I) and male (Fig 5J) NAPE-PLD KO mice [Fig. 5I, Female: Interaction: F (2, 18) = 39.76, P<0.0001; Condition: F (2, 18) = 289.4, P<0.0001; Genotype: F (1, 18) = 0.1047, P=0.7500. Fig. 5J, Male: Interaction: F (2, 18) = 14.75, P=0.0002; Condition: F (2, 18) = 78.56, P<0.0001; Genotype F (1, 18) = 2.330, P=0.1443]. c-Fos+ cell counts correlated positively in both WT (Fig 5K) and NAPE-PLD KO (Fig 5L) mice exposed to water and TMT, though the slope was flatter in NAPE-PLD KO mice [WT: Spearman r: 0.7500, P=0.0012; NAPE-PLD KO: Spearman r: 0.6441, P=0.0085].

### NAPE-PLD KO mice exhibit context-dependent and sexually dimorphic alterations in circulating corticosterone

For further mechanistic understanding of how NAPE-PLD deletion would affect HPA-axis functionality in context of the observed behavioral phenotype, a subset of WT and NAPE-PLD KO mice were sacrificed immediately after odor exposure and cardiac blood was collected for CORT quantification. For how the context of testing conditions (placement in a fume hood) would affect responses, we also included behaviorally naïve subjects into our analyses as performed with immunohistochemical analyses (Fig 6A). CORT levels were modestly higher in experimentally naïve NAPE-PLD KO mice compared to WT controls (Fig 6B) [Interaction: F (1, 30) = 2.651, P=0.1139; Sex: F (1, 30) = 2.804, P=0.1044; Genotype: F (1, 30) = 6.635, P=0.0152]. CORT levels increased in WT (Fig 6C) and NAPE-PLD KO (Fig 6D) mice as context aversiveness increased [Fig 6C, WT: Interaction: F (2, 44) = 1.038, P=0.3628; Context: F (2, 44) = 22.04, P<0.0001; Sex: F (1, 44) = 2.291, P=0.1373; Fig 6D, NAPE-PLD KO: Interaction: F (2, 37) = 1.830, P=0.1746; Context: F (2, 37) = 28.66, P<0.0001; Sex: F (1, 37) = 0.1269, P=0.7237]. While female NAPE-PLD KO mice only trended toward increased CORT levels across contexts compared to female WT mice (Fig 6E), male NAPE-PLD KO mice displayed persistent elevation of CORT levels, especially in response to TMT exposure [Fig 6E, Females: Interaction: F (2, 40) = 0.8350, P=0.4413; Context: F (2, 40) = 19.06, P<0.0001; Genotype: F (1, 40) = 3.082, P=0.0868; 6F, Males: Interaction: F (2, 41) = 5.200, P=0.0097; Context: F (2, 41) = 36.76, P<0.0001; Genotype: F (1, 41) = 18.32, P=0.0001].

**Figure 6.**
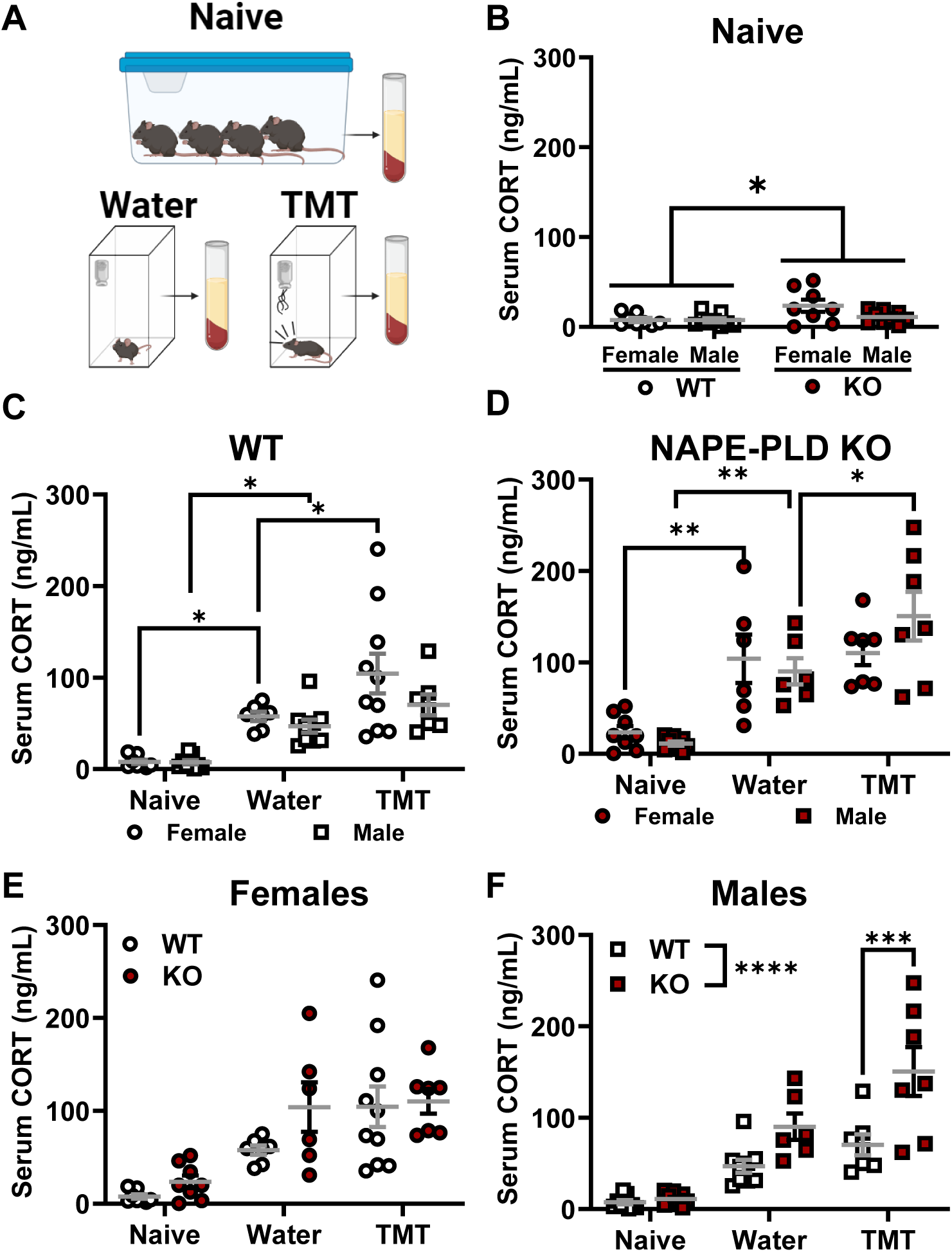
Circulating corticosterone levels were elevated among NAPE-PLD KO mice in a sexually dimorphic manner. Serum samples were obtained from experimentally naïve mice, as well as immediately after water or TMT exposure in the fume hood to determine levels of circulating corticosterone (CORT) as seen in experimental schematic **A**. Compared to experimentally naïve WT mice, NAPE-PLD KO mice exhibited a modest but statistically significant increase in levels of circulating CORT **(B)**. In WT **(C)** and NAPE-PLD KO **(D)** mice, levels of circulating CORT increased as a function of context aversiveness in both sexes. CORT levels in female NAPE-PLD KO mice did not differ from female WT CORT levels across contexts **(E)**. In contrast, CORT levels were elevated in male NAPE-PLD KO mice compared to WT controls, especially after TMT exposure **(F)**. Data are presented as the mean ± S.E.M. Multiple comparisons from two-way ANOVA followed by Bonferroni post-hoc test: ****p<0.0001, ***p<0.001, **p<0.01, * p<0.05 for shown comparisons. Sample size: n=8-12 per sex per genotype per odor condition.

## Discussion

CB1 signaling and NAE degradation (via FAAH) are known to modulate stress responsiveness (Haller et al., 2004, 2009). To our knowledge, however, this is the first study to describe the *context-dependent* nature in which NAE synthesis via NAPE-PLD may regulate reactivity to stress. Biochemical consensus now supports a role for NAPE-PLD as the primary enzyme involved in NAE formation (Leishman et al., 2016). Initial *in vitro* studies suggested that NAPE-PLD catalyzed the formation of AEA (Di Marzo et al., 1994; Okamoto et al., 2005). However, brain AEA levels from the first NAPE-PLD knockout (KO) mouse line (known in the literature as the Cravatt line), did not differ from wild type controls, a finding which led to identification of alternative pathways of AEA formation (Leung et al., 2006; Liu et al., 2008). A recent behavioral characterization of the Cravatt NAPE-PLD KO line demonstrated that deletion of NAPE-PLD had little impact on baseline affective behaviors (Chen et al., 2023). Subsequent knockout (KO) mouse lines that removed exon 3 (later found to be the main catalytic site) instead of exon 4 (removed in the Cravatt line) exhibited dramatic reduction in levels of AEA as well as other NAEs (Liu et al., 2008; Tsuboi et al., 2011; Leishman et al., 2016).

In line with a recent report using the Cravatt line (Chen et al., 2023), we found only a few behavioral differences in group-housed mice in the open field and the light-dark box tests. In contrast to the heightened level of exploratory rearing in the open field test observed in FAAH KO mice (Bambico et al., 2010), male NAPE-PLD KO mice in our study spent less time in exploratory rearing behaviors than WT counterparts. NAPE-PLD KO mice only began to exhibit a stronger anxiety-like phenotype after 3 weeks of single housing, while WT mice were relatively unaffected by this housing manipulation in our hands. This is potentially of interest given previous documentations of aberrant AEA signaling produced by lengthier post-weaning social isolation in rats (Malone et al., 2008; Robinson et al., 2010; Sciolino et al., 2010; Carnevali et al., 2020). The anxiety-like phenotype was especially pronounced amongst aged (35+ weeks old) single-housed NAPE-PLD KO mice. CB1 KO mice exhibit an anxiety-like behavioral phenotype that is attenuated if mice are habituated to the testing environment and tested under low-light (i.e., less aversive) conditions (Haller et al., 2004). Given that CB1 KO mice exhibit age-dependent dysregulation of anxiety-like behaviors and AEA metabolism (Maccarrone et al., 2002), NAPE-PLD could also potentially regulate stress coping differentially during development. Further work is required to fully examine this hypothesis.

c-Fos is an intermediate-early gene whose expression is often used as a nonspecific marker of neuronal activity, as its expression increases in response to large influxes of calcium (Chung, 2015). In our study, NAPE-PLD KO mice exhibited a basal reduction in c-Fos staining within both the CA3 region and dentate gyrus (DG) of the hippocampus. The reduced number of c-Fos-positive neurons observed in experimentally naïve baseline conditions raises the possibility that NAPE-PLD activity during development may regulate the maturation of brain circuits and diminish the number of neurons involved in a given population activity pattern (Berghuis et al., 2007). While animals were not challenged in their home cage environment, it is important to emphasize the differences in the behavioral tests were only observed in conditions that were stressful for the animals. Thus, the absence of NAPE-PLD during embryonic development may lead to an altered threshold for the recruitment of neuronal ensembles encoding context-dependent (novel environment) or innate (fox odor) stress and fear responses may partly explain the complex behavioral effects found the NAPE-PLD global KO mice in the present study. Recently, activity of NAPE-PLD in specific neuronal populations was shown to potentially supports stress adaptation in a chronic social defeat paradigm (Tevosian et al., 2023).

For example, in response to water exposure (no odor), c-Fos expression and behavioral freezing was amplified in NAPE-PLD KO mice, potentially due to the aversiveness of the fume hood, whose ambient sound level was at about 77 dB. The higher behavioral and neuronal sensitivity to the testing environment may have masked the consequences of TMT exposure, resulting in an elevation of both behavioral and neurochemical responses to a lesser extent in the NAPE-PLD KO mice compared to WT mice. NAPE-PLD KO mice thus exhibited a blunted response to TMT at the behavioral and neuronal level. While it is possible that olfaction could be somehow impaired by NAPE-PLD deletion, it is also possible that NAPE-PLD’s activity facilitates the dynamic range, or flexibility, of the HPA axis, a hypothesis which could be elucidated in future studies.

In NAPE-PLD KO mice, we observed sexual dimorphism in dysregulation of corticosterone levels in response to stress. Given that we didn’t observe reliable sex differences in c-Fos staining within the PVN in any stress condition, the discrepancy between c-Fos staining and corticosterone levels could indicate that other feedback mechanisms within the HPA axis are affected in a sexually dimorphic manner by NAPE-PLD deletion (CORT breakdown, pituitary function, etc.). Sexual dimorphism in HPA axis functionality is well-documented (Goel et al., 2014; Heck and Handa, 2019), and NAE levels in humans and rodents differ both as a function of biological sex (male vs female) and as a function of estrous/menstrual cycle in rodents and humans (Bradshaw et al., 2006; El-Talatini et al., 2010; Cui et al., 2017). Interestingly, pharmacological manipulations of both CB1 receptors and FAAH produce cycle-dependent effects in rodents (Kim et al., 2022; Salemme et al., 2023) To our knowledge, this is the first study to suggest that NAPE-PLD regulates the HPA axis differently in males and females. While we did not examine stage of estrous cycle in females or perform manipulations of sex hormones (gonadectomy), these finding could set a precedent for future investigations regarding NAPE-PLD and sex differences in the HPA axis.

The enzyme fatty acid amide hydrolase (FAAH) is well known to catalyze AEA degradation (for review, see Deutsch, 2016) and is implicated in stress processing. To date, investigations into the role of AEA in stress responses have almost exclusively focused on increasing AEA levels by inhibiting FAAH (Lutz et al., 2015). Thus, it is noteworthy that FAAH knockout (KO) mice, which exhibit elevated levels of AEA, also show reductions in anxiety-like behavior compared to wildtype (WT) controls (Bambico et al., 2010). In healthy individuals, acute stress can increase blood levels of AEA and NAEs. In patients with PTSD or depression, AEA levels often negatively correlate with cortisol levels and/or symptoms in patients with PTSD or depression (Dlugos et al., 2012; Hill et al., 2013; Trevino et al., 2022; Walther et al., 2023). FAAH inhibitors are currently under evaluation as a potential treatment for anxiety and post-traumatic stress disorders (Paulus et al., 2021). Studies of a single-nucleotide polymorphism (SNP) in FAAH (C385A), which reduces its activity and thereby raises AEA levels, also link this SNP to elevated AEA levels and improved fear extinction in both mice and humans (Dincheva et al., 2015; Mayo et al., 2018; Spohrs et al., 2022). Both dysregulation of the HPA axis as well AEA levels have also been documented in autistic individuals (Karhson et al., 2018; Makris et al., 2022) as well as individuals with schizophrenia (Giuffrida et al., 2004; Minichino et al., 2019). Interestingly, a recent study found that individuals with obsessive-compulsive disorder (OCD) exhibited a downregulation of NAPE-PLD gene expression when compared to healthy controls (Bellia et al., 2024). These observations link dysregulation of AEA, and potentially NAPE-PLD, to stress and neuropsychiatric indications. NAE signaling is complicated, as the products of NAPE-PLD-mediated metabolism potentially exert effects on various targets include CB1, CB2, peroxisome proliferator receptor-α (PPARα), and transient receptor potential vanilloid cation channel family subtype V member 1 (TRPV1) among others (Mock et al., 2023).

Understanding the nuances of how NAEs modulate HPA axis function could open new doors for diagnosis/therapies for stress disorders. In early explorations if the FAAH inhibitor URB597, it was that elevating NAE levels only produced anxiolytic effects under more aversive testing conditions (Haller et al., 2009). This is of interest, given that in the present study, NAPE-PLD KO mice (with lower NAE levels) exhibited alterations in response to stress at a behavioral, neuronal, and hormonal level that were dependent upon context aversiveness. Further investigation using cell-dependent invalidation is warranted to further dissect the molecular and cellular mechanism by which NAPE-PLD controls the HPA axis. Given the precedent that SNPs in FAAH can potentially bestow increased resilience to stress (Dincheva et al., 2015; Spohrs et al., 2022), SNPs in NAPE-PLD could potentially serve as a risk factor for the development of stress disorders. Together, our results support the likely significance in NAPE-PLD’s activity as a regulator of stress responses.

## Acknowledgements

This work was supported by DA047858 (to AGH and KM), DA009158 (to AGH). TW was supported National Institute on Drugs Abuse T32 training grant DA024628, the Harlan Scholars Research Program, and Gill Graduate Research Fellowship. Experimental schematics in figures were created with BioRender.com.

## Author Contributions

TJW, DD, EFS, SS, and FK performed research and analyzed data. TW, DD, and AGH designed experiments. AGH and IK oversaw the project. TJW, DD, and AGH wrote the manuscript. AGH and KM generated resources that supported the project.

## References

Anisimova M, Lamothe-Molina PJ, Franzelin A, Aberra AS, Hoppa MB, Gee CE, Oertner TG (2023) Neuronal FOS reports synchronized activity of presynaptic neurons. bioRxiv Available at: 10.1101/2023.09.04.556168 [Accessed November 29, 2023].

Bambico FR, Cassano T, Dominguez-Lopez S, Katz N, Walker CD, Piomelli D, Gobbi G (2010) Genetic deletion of fatty acid amide hydrolase alters emotional behavior and serotonergic transmission in the dorsal raphe, prefrontal cortex, and hippocampus. Neuropsychopharmacology 35:2083–2100.

Bankhead P, Loughrey MB, Fernández JA, Dombrowski Y, Mcart DG, Dunne PD, Mcquaid S, Gray RT, Murray LJ, Coleman HG, James JA, Salto-Tellez M, Hamilton PW (2017) QuPath: Open source software for digital pathology image analysis. Sci Rep 7:16878 Available at: www.nature.com/scientificreports/ [Accessed August 31, 2023].

Bellia F, Girella A, Annunzi E, Benatti B, Vismara M, Priori A, Festucci F, Fanti F, Compagnone D, Adriani W, Dell’Osso B, D’Addario C (2024) Selective alterations of endocannabinoid system genes expression in obsessive compulsive disorder. Translational Psychiatry 2024 14:1 14:1–10 Available at: https://www.nature.com/articles/s41398-024-02829-8 [Accessed April 23, 2024].

Berghuis P, Rajnicek A, Morozov Y, Ross R, Mulder J, Urbán G, Monory K, Marsicano G, Matteoli M, Canty A, Irving A, Katona I, Yanagawa Y, Rakic P, Lutz B, Mackie K, Harkany T (2007) Hardwiring the brain: endocannabinoids shape neuronal connectivity. Science (1979) 316:2061–2065.

Bradshaw HB, Rimmerman N, Krey JF, Walker JM (2006) Sex and hormonal cycle differences in rat brain levels of pain-related cannabimimetic lipid mediators. Am J Physiol Regul Integr Comp Physiol 291:349–358 Available at: https://journals.physiology.org/doi/10.1152/ajpregu.00933.2005 [Accessed August 31, 2023].

Carnevali L, Statello R, Vacondio F, Ferlenghi F, Spadoni G, Rivara S, Mor M, Sgoifo A (2020) Antidepressant-like effects of pharmacological inhibition of FAAH activity in socially isolated female rats. Eur Neuropsychopharmacol 32:77–87 Available at: https://pubmed.ncbi.nlm.nih.gov/31948828/ [Accessed September 1, 2023].

Chávez AE, Chiu CQ, Castillo PE (2010) TRPV1 activation by endogenous anandamide triggers postsynaptic LTD in dentate gyrus. Nat Neurosci 13:1511 Available at: /pmc/articles/PMC3058928/ [Accessed September 2, 2023].

Chen I, Murdaugh LB, Miliano C, Dong Y, Gregus AM, Buczynski MW (2023) NAPE-PLD regulates specific baseline affective behaviors but is dispensable for inflammatory hyperalgesia. Available at: 10.1016/j.ynpai.2023.100135 [Accessed August 30, 2023].

Chung L (2015) A Brief Introduction to the Transduction of Neural Activity into Fos Signal. Dev Reprod 19:61 Available at: /pmc/articles/PMC4801051/ [Accessed September 1, 2023].

Cui N, Wang L, Wang W, Zhang J, Xu Y, Jiang L, Hao G (2017) The correlation of anandamide with gonadotrophin and sex steroid hormones during the menstrual cycle The correlation of anandamide with gonadotrophin and sex steroid hormones during the menstrual cycle in Asian women. Iran J Basic Med Sci 20:1268–1274.

Deutsch DG (2016) A Personal Retrospective: Elevating Anandamide (AEA) by Targeting Fatty Acid Amide Hydrolase (FAAH) and the Fatty Acid Binding Proteins (FABPs). Frontiers in Pharmacology | www.frontiersin.org 7 Available at: www.frontiersin.org [Accessed August 29, 2023].

Di Marzo V, Fontana A, Cadas H, Schinelli S, Cimino G, Schwartz J-C, Piomelli D (1994) Formation and inactivation of endogenous cannabinoid anandamide in central neurons. Nature 372:686–691.

Dincheva I, Drysdale AT, Hartley CA, Johnson DC, Jing D, King EC, Ra S, Gray JM, Yang R, DeGruccio AM, Huang C, Cravatt BF, Glatt CE, Hill MN, Casey BJ, Lee FS (2015) FAAH genetic variation enhances fronto-amygdala function in mouse and human. Nat Commun 6:6395 Available at: /pmc/articles/PMC4351757/ [Accessed August 29, 2023].

Dlugos A, Childs E, Stuhr KL, Hillard CJ, De Wit H (2012) Acute Stress Increases Circulating Anandamide and Other N-Acylethanolamines in Healthy Humans. Neuropsychopharmacology 37:2416–2427 Available at: www.neuropsychopharmacology.org [Accessed August 31, 2023].

Dvorakova M, Kubik-Zahorodna A, Straiker A, Sedlacek R, Hajkova A, Mackie K, Blahos J (2021) SGIP1 is involved in regulation of emotionality, mood, and nociception and modulates in vivo signalling of cannabinoid CB1 receptors. Br J Pharmacol 178:1588– 1604.

Egertová M, Simon GM, Cravatt BF, Elphick MR (2008) Localization of N-acyl phosphatidylethanolamine phospholipase D (NAPE-PLD) expression in mouse brain: A new perspective on N-acylethanolamines as neural signaling molecules. Journal of Comparative Neurology 506:604–615.

El-Talatini MR, Taylor AH, Konje JC (2010) The relationship between plasma levels of the endocannabinoid, anandamide, sex steroids, and gonadotrophins during the menstrual cycle. Fertil Steril 93:1989–1996.

Giuffrida A, Leweke M, Gerth CW, Schreiber D, Koethe D, Faulhaber J, Klosterkö Tter J, Piomelli D (2004) Cerebrospinal Anandamide Levels are Elevated in Acute Schizophrenia and are Inversely Correlated with Psychotic Symptoms. Neuropsychopharmacology 29:2108–2114 Available at: http://www.acnp.org/citations/ [Accessed September 1, 2023].

Goel N, Workman JL, Lee TT, Innala L, Viau V (2014) Sex Differences in the HPA Axis. Compr Physiol 4:1121–1155 Available at: https://onlinelibrary.wiley.com/doi/full/10.1002/cphy.c130054 [Accessed August 31, 2023].

Grueter BA, Brasnjo G, Malenka RC (2010) Postsynaptic TRPV1 triggers cell type-specific long-term depression in the nucleus accumbens. Nat Neurosci 13:1519–1526.

Haller J, Barna I, Barsvari B, Gyimesi Pelczer K, Yasar S, Panlilio L V, Goldberg S (2009) Interactions between environmental aversiveness and the anxiolytic effects of enhanced cannabinoid signaling by FAAH inhibition in rats. Psychopharmacology (Berl) 204:607–616.

Haller J, Varga B, Ledent C, Barna I, Freund TF (2004) Context-dependent effects of CB1 cannabinoid gene disruption on anxiety-like and social behaviour in mice. European Journal of Neuroscience 19:1906–1912 Available at: https://onlinelibrary.wiley.com/doi/full/10.1111/j.1460-9568.2004.03293.x [Accessed August 30, 2023].

Heck AL, Handa RJ (2019) Sex differences in the hypothalamic–pituitary–adrenal axis’ response to stress: an important role for gonadal hormones. Neuropsychopharmacology 44:45–58.

Hill MN, Bierer LM, Makotkine I, Golier JA, Galea S, McEwen BS, Hillard CJ, Yehuda R (2013) Reductions in circulating endocannabinoid levels in individuals with post-traumatic stress disorder following exposure to the world trade center attacks. Psychoneuroendocrinology 38:2952–2961.

Hillard CJ, Beatka M, Sarvaideo J (2017) Endocannabinoid Signaling and the Hypothalamic-Pituitary-Adrenal Axis Introduction to the Components and Paradigms of CNS Endocannabinoid Signaling. Compr Physiol 7:1–15 Available at: https://onlinelibrary.wiley.com/doi/10.1002/cphy.c160005 [Accessed September 1, 2023].

Iyer V, Saberi SA, Pacheco R, Sizemore EF, Stockman S, Kulkarni A, Cantwell L, Thakur GA, Hohmann AG (2024) Negative allosteric modulation of CB1 cannabinoid receptor signaling suppresses opioid-mediated tolerance and withdrawal without blocking opioid antinociception. Neuropharmacology 257.

Janitzky K, D’Hanis W, Kröber A, Schwegler H (2015) TMT predator odor activated neural circuit in C57BL/6J mice indicates TMT-stress as a suitable model for uncontrollable intense stress. Brain Res 1599:1–8.

Karhson DS, Krasinska KM, Dallaire JA, Libove RA, Phillips JM, Chien AS, Garner JP, Hardan AY, Parker KJ (2018) Plasma anandamide concentrations are lower in children with autism spectrum disorder. Mol Autism 9 Available at: /pmc/articles/PMC5848550/ [Accessed November 25, 2021].

Kim HJJ, Zagzoog A, Black T, Baccetto SL, Ezeaka UC, Laprairie RB (2022) Impact of the mouse estrus cycle on cannabinoid receptor agonist-induced molecular and behavioral outcomes. Pharmacol Res Perspect 10:e00950 Available at: https://onlinelibrary.wiley.com/doi/full/10.1002/prp2.950 [Accessed August 31, 2023].

Leishman E, Mackie K, Luquet S, Bradshaw HB (2016) Lipidomics profile of a NAPE-PLD KO mouse provides evidence of a broader role of this enzyme in lipid metabolism in the brain. Biochim Biophys Acta Mol Cell Biol Lipids 1861:491–500.

Leung D, Saghatelian A, Simon GM, Cravatt BF (2006) Inactivation of N-Acyl Phosphatidylethanolamine Phospholipase D Reveals Multiple Mechanisms for the Biosynthesis of Endocannabinoids. Available at: http://pubs.acs.org.

Liu J, Wang L, Harvey-White J, Huang BX, Kim H-Y, Luquet S, Palmiter RD, Krystal G, Rai R, Mahadevan A, Razdan RK, Kunos G (2008) Multiple Pathways Involved in the Biosynthesis of Anandamide. Neuropharmacology 54:1–7.

Lutz B, Marsicano G, Maldonado R, Hillard CJ (2015) The endocannabinoid system in guarding against fear, anxiety and stress. Nat Rev Neurosci 16:705–718.

Maccarrone M, Valverde O, Barbaccia ML, Castan A, Maldonado R, Ledent C, Parmentier M, Finazzi-Agro A (2002) Age-related changes of anandamide metabolism in CB 1 cannabinoid receptor knockout mice: correlation with behaviour. European Journal of Neuroscience 15:1779–1186.

Maciel IDS, Abreu GHDD, Johnson CT, Bonday R, Bradshaw HB, Mackie K, Lu HC (2022) Perinatal CBD or THC Exposure Results in Lasting Resistance to Fluoxetine in the Forced Swim Test: Reversal by Fatty Acid Amide Hydrolase Inhibition. Cannabis Cannabinoid Res 7:318–327.

Makris G, Agorastos A, Chrousos GP, Pervanidou P (2022) Stress System Activation in Children and Adolescents With Autism Spectrum Disorder. Front Neurosci 15.

Malone DT, Kearn CS, Chongue L, Mackie K, Taylor DA (2008) Effect of social isolation on CB1 and D2 receptor and fatty acid amide hydrolase expression in rats. Neuroscience 152:265– 272.

Mayo LM, Asratian A, Lindé J, Holm L, Nätt D, Augier G, Stensson N, Vecchiarelli HA, Balsevich G, Aukema RJ, Ghafouri B, Spagnolo PA, Lee FS, Hill MN, Heilig M (2018) Protective effects of elevated anandamide on stress and fear-related behaviors: translational evidence from humans and mice. Molecular Psychiatry 2018 25:5 25:993– 1005 Available at: https://www.nature.com/articles/s41380-018-0215-1 [Accessed August 29, 2023].

Minichino A, Senior M, Brondino N, Zhang SH, Godwlewska BR, Burnet PWJ, Cipriani A, Lennox BR (2019) Measuring Disturbance of the Endocannabinoid System in Psychosis: A Systematic Review and Meta-analysis. JAMA Psychiatry 76:914–923 Available at: https://pubmed.ncbi.nlm.nih.gov/31166595/ [Accessed September 1, 2023].

Mock ED, Gagestein B, van der Stelt M (2023) Anandamide and other N-acylethanolamines: A class of signaling lipids with therapeutic opportunities. Prog Lipid Res 89:101194.

Murphy M, Mills S, Winstone J, Leishman E, Wager-Miller J, Bradshaw H, Mackie K (2017) Chronic Adolescent Δ9-Tetrahydrocannabinol Treatment of Male Mice Leads to Long-Term Cognitive and Behavioral Dysfunction, Which Are Prevented by Concurrent Cannabidiol Treatment. Cannabis Cannabinoid Res 2:235–246.

Nyilas R, Dudok B, Urbán GM, Mackie K, Watanabe M, Cravatt BF, Freund TF, Katona I (2008) Enzymatic machinery for endocannabinoid biosynthesis associated with calcium stores in glutamatergic axon terminals. Journal of Neuroscience 28:1058–1063.

Okamoto Y, Morishita J, Wang J, Schmid PC, Krebsbach RJ, Schmid HHO, Ueda N (2005) Mammalian cells stably overexpressing N-acylphosphatidylethanolamine-hydrolysing phospholipase D exhibit significantly decreased levels of N-acylphosphatidylethanolamines. Biochemical Journal 389:241 Available at: /pmc/articles/PMC1184557/ [Accessed August 29, 2023].

Paulus MP, Stein MB, Simmons AN, Risbrough VB, Halter R, Chaplan SR (2021) The effects of FAAH inhibition on the neural basis of anxiety-related processing in healthy male subjects: a randomized clinical trial. Neuropsychopharmacology 46:1011–1019.

Pennington ZT, Diego KS, Francisco TR, LaBanca AR, Lamsifer SI, Liobimova O, Shuman T, Cai DJ (2021) ezTrack—A Step-by-Step Guide to Behavior Tracking. Curr Protoc 1.

Pennington ZT, Dong Z, Feng Y, Vetere LM, Page-Harley L, Shuman T, Cai DJ (2019) ezTrack: An open-source video analysis pipeline for the investigation of animal behavior. Scientific Reports 2019 9:1 9:1–11 Available at: https://www.nature.com/articles/s41598-019-56408-9 [Accessed August 31, 2023].

Pickel VM, Shobin ET, Lane DA, Mackie K (2012) Cannabinoid-1 receptors in the mouse ventral pallidum are targeted to axonal profiles expressing functionally opposed opioid peptides and contacting N-acylphosphatidylethanolamine-hydrolyzing phospholipase D terminals. Neuroscience 227:10–21 Available at: http://www.ibroneuroscience.org/article/S0306452212007750/fulltext [Accessed September 2, 2023].

Puente N, Cui Y, Lassalle O, Lafourcade M, Georges F, Venance L, Grandes P, Manzoni OJ (2011) Polymodal activation of the endocannabinoid system in the extended amygdala. Nature Neuroscience 2011 14:12 14:1542–1547 Available at: https://www.nature.com/articles/nn.2974 [Accessed September 2, 2023].

Reguero L, Puente N, Elezgarai I, Ramos-Uriarte A, Gerrikagoitia I, Bueno-López JL, Doñate F, Grandes P (2014) Subcellular localization of NAPE-PLD and DAGL-α in the ventromedial nucleus of the hypothalamus by a preembedding immunogold method. Histochem Cell Biol 141:543–550.

Robinson SA, Loiacono RE, Christopoulos A, Sexton PM, Malone DT (2010) The effect of social isolation on rat brain expression of genes associated with endocannabinoid signaling. Brain Res 1343:153–167.

Rosen JB, Asok A, Chakraborty T (2015) The smell of fear: Innate threat of 2,5-dihydro-2,4,5-trimethylthiazoline, a single molecule component of a predator odor. Front Neurosci 9:1–12.

Salemme BW, Raymundi AM, Sohn JMB, Stern CA (2023) The Estrous Cycle Influences the Effects of Fatty Acid Amide Hydrolase and Monoacylglycerol Lipase Inhibition in the Anxiety-Like Behavior in Rats. Cannabis Cannabinoid Res Available at: https://www.liebertpub.com/doi/10.1089/can.2022.0329 [Accessed August 31, 2023].

Sciolino NR, Bortolato M, Eisenstein SA, Fu J, Oveisi F, Hohmann AG, Piomelli D (2010) Social isolation and chronic handling alter endocannabinoid signaling and behavioral reactivity to context in adult rats. Neuroscience 168:371–386 Available at: http://rsb.info.nih.gov/ [Accessed September 1, 2023].

Slivicki RA, Saberi SA, Iyer V, Vemuri VK, Makriyannis A, Hohmann AG (2018) Brain-permeant and -impermeant inhibitors of fatty acid amide hydrolase synergize with the opioid analgesic morphine to suppress chemotherapy-induced neuropathic nociception without enhancing effects of morphine on gastrointestinal transit. Journal of Pharmacology and Experimental Therapeutics 367:551–563.

Spohrs J, Ulrich M, Grön G, Plener PL, Abler B (2022) FAAH polymorphism (rs324420) modulates extinction recall in healthy humans: an fMRI study. Eur Arch Psychiatry Clin Neurosci 272:1495–1504 Available at: 10.1007/s00406-021-01367-4 [Accessed August 29, 2023].

Tan SY, Yip A (2018) Hans Selye (1907–1982): Founder of the stress theory. Singapore Med J 59:170–171 Available at: 10.11622/smedj.2018043 [Accessed August 31, 2023].

Tevosian M, Todorov H, Lomazzo E, Bindila L, Ueda N, Bassetti D, Warm D, Kirischuk S, Luhmann HJ, Gerber S, Lutz B (2023) NAPE-PLD deletion in stress-TRAPed neurons results in an anxiogenic phenotype. Transl Psychiatry 13:152 Available at: 10.1038/s41398-023-02448-9 [Accessed August 30, 2023].

Trevino CM, Hillard CJ, Szabo A, Deroon-Cassini TA (2022) Serum Concentrations of the Endocannabinoid, 2-Arachidonoylglycerol, in the Peri-Trauma Period Are Positively Associated with Chronic Pain Months Later. Biomedicines 10.

Tsuboi K, Okamoto Y, Ikematsu N, Inoue M, Shimizu Y, Uyama T, Wang J, Deutsch DG, Burns MP, Ulloa NM, Tokumura A, Ueda N (2011) Enzymatic formation of N-acylethanolamines from N-acylethanolamine plasmalogen through N-acylphosphatidylethanolamine-hydrolyzing phospholipase D-dependent and -independent pathways. Biochimica et Biophysica Acta (BBA) -Molecular and Cell Biology of Lipids 1811:565–577.

Walther A, Kirschbaum C, Wehrli S, Rothe N, Penz M, Wekenborg M, Gao W (2023) Depressive symptoms are negatively associated with hair N-arachidonoylethanolamine (anandamide) levels: A cross-lagged panel analysis of four annual assessment waves examining hair endocannabinoids and cortisol. Prog Neuropsychopharmacol Biol Psychiatry 121:278–5846 Available at: 10.1016/j.pnpbp.2022.110658 [Accessed August 31, 2023].

